# Detection of human adaptation during the past 2,000 years

**DOI:** 10.1101/052084

**Authors:** Yair Field, Evan A Boyle, Natalie Telis, Ziyue Gao, Kyle J. Gaulton, David Golan, Loic Yengo, Ghislain Rocheleau, Philippe Froguel, Mark I. McCarthy, Jonathan K. Pritchard

## Abstract

Detection of recent natural selection is a challenging problem in population genetics, as standard methods generally integrate over long timescales. Here we introduce the Singleton Density Score (SDS), a powerful measure to infer very recent changes in allele frequencies from contemporary genome sequences. When applied to data from the UK10K Project, SDS reflects allele frequency changes in the ancestors of modern Britons during the past 2,000 years. We see strong signals of selection at lactase and HLA, and in favor of blond hair and blue eyes. Turning to signals of polygenic adaptation we find, remarkably, that recent selection for increased height has driven allele frequency shifts across most of the genome. Moreover, we report suggestive new evidence for polygenic shifts affecting many other complex traits. Our results suggest that polygenic adaptation has played a pervasive role in shaping genotypic and phenotypic variation in modern humans.

## Main Text

Understanding the genetic basis of adaptation is a central goal in evolutionary biology. Most work in humans and other species has focused on identifying signals of strong selection at individual loci [1–3]. In humans, this work has identified loci involved in adaptations to diet, altitude, and disease resistance, and toward lighter pigmentation in northern populations [4 –8].

Early approaches to identifying signals of selection focused on detecting signals of “hard sweeps”, in which new mutations are immediately favored and sweep through a population [9 –12]. But many of the clearest examples of selection involve recent adaptive changes in allele frequencies at pre-existing variants. These so-called “soft sweeps” [13,14] may occur when an existing variant becomes favored due to environmental change, or when a population colonizes a new location [6,15]. Further, it has been hypothesized that another main mechanism of adaptation is through polygenic adaptation, a rapid process in which small allele frequency changes at existing variants across the genome drive adaptation of complex traits [16]. The clearest example of polygenic adaptation is for height in Europeans [17 –19].

However, it is difficult to measure very recent changes in allele frequencies. Methods designed to detect recent hard sweeps integrate information over the entire history of a sample, and effectively measure selection over long timescales—typically 25,000 years or more [20]. An alternative widely-used approach is to test for differences in allele frequencies between extant populations [6,21,22]. Such methods can be powerful for detecting selection on standing variation, but generally rely on the availability of closely related populations with divergent selective pressures. Suitable population pairs may not exist, and if they do, it may be unclear which selection pressures differ between them. Alternatively, recent papers have compared allele frequencies from ancient DNA with modern samples [23,24]. Use of ancient DNA is potentially powerful but relies on the availability of large numbers of well-preserved ancestral samples.

Here we introduce a new technique for measuring very recent changes in allele frequencies. Our measure, the Singleton Density Score (SDS), uses modern whole-genome sequence data, and will thus be widely applicable. SDS focuses on patterns of variation around each SNP to infer recent changes in the relative frequencies of the two alleles. The key idea is that recent selection distorts the ancestral genealogy of sampled haplotypes at a selected site. In particular, the terminal (tip) branches of the genealogy tend to be shorter for the favored allele than for the disfavored allele, and hence, haplotypes carrying the favored allele will tend to carry fewer singleton mutations (Fig. 1A-C and SOM).

**Figure 1.**
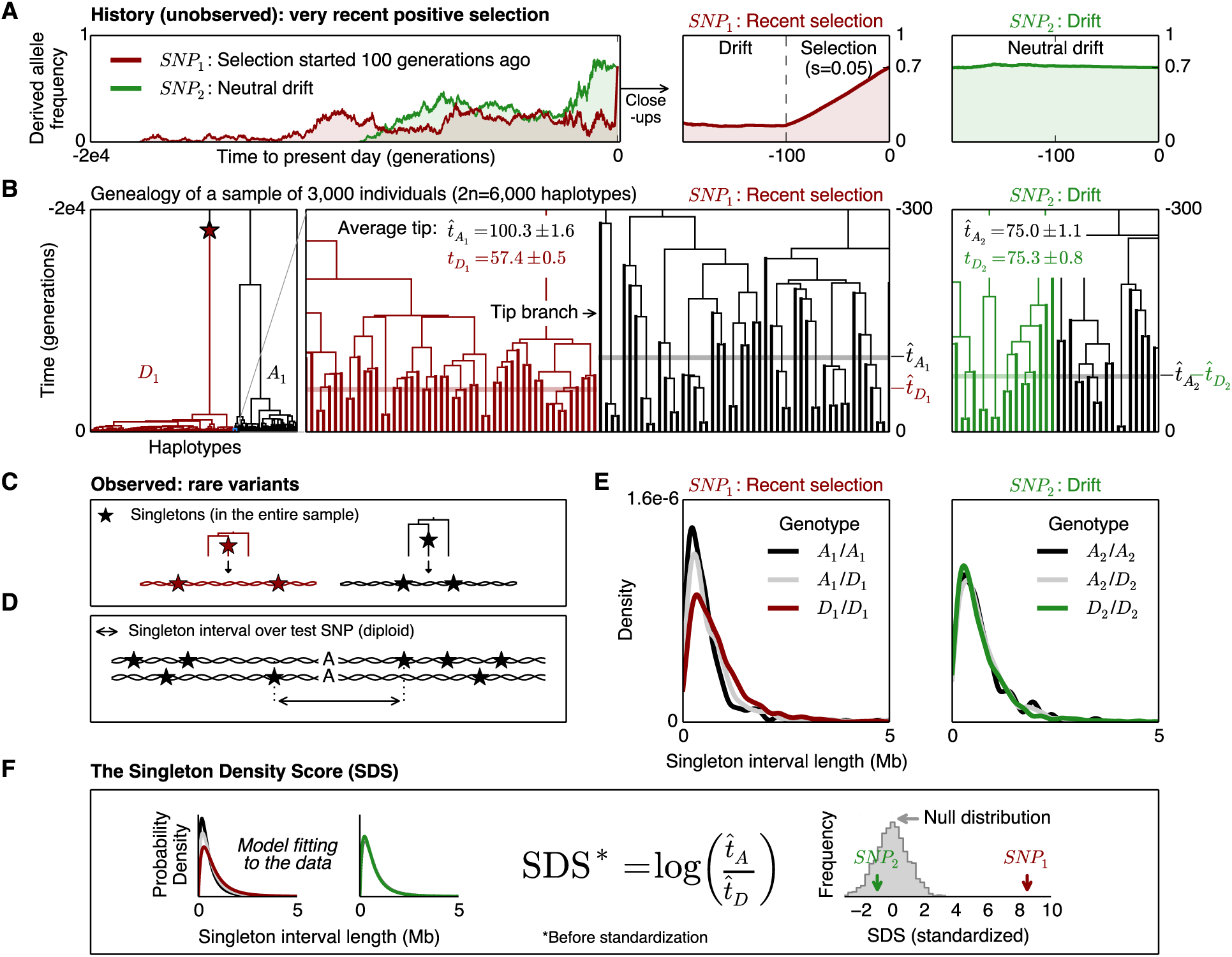
Illustration of the SDS method. ***A.** Simulated frequency trajectories for derived alleles at two different SNPs, each at frequency 0.7 in the present day. The allele marked in green was selectively neutral at all times (left and right panels). The red allele was neutral until 100 generations before the present, at which point it became favored and spread rapidly to its present frequency (left and middle panels). **B.** The corresponding genealogy of a sample of 3,000 diploids for the selected SNP (left and middle panels). Lineages carrying the derived allele (D1) are in red; ancestral lineages (A1) are in black. Blow-up of a small part of the genealogy (middle panel) illustrates that tip branches carrying the favored allele (red) are shorter on average than those carrying the disfavored allele (black). Blow-up of part of the genealogy for the neutral SNP (right panel) shows no significant difference in average tip length between the two alleles. **C.** Because favored alleles (red) tend to have shorter tips, their haplotypes tend to have lower singleton density. **D.** For each individual we compute the total distance between nearest singletons in each direction from the test SNP. **E.** Distribution of singleton distances as a function of genotype at the test SNP for the selected site (left) and neutral site (right). **F.** Mean tip length t is estimated for each allele from a likelihood model. Unstandardized SDS is a log ratio of estimated tip lengths; this is standardized to mean 0, variance 1 within 1% bins of derived allele frequency. The resulting distribution is standard normal under the null hypothesis (SOM). In this simulated example, the neutral site is not significant (p=0.35) while the selected site is highly significant (p=1×10^−17^ in favor of the derived allele)*.

To capture this effect, we use the sum of distances to the nearest singleton in each direction from a test SNP as a summary statistic (Fig. 1D). Since singletons are generally unphased in current data, the singleton distances are computed from unphased diploid data. We then use the distributions of distances for each of the three genotypes at the test SNP to compute a maximum likelihood estimate of the log ratio of mean tip-branch lengths for the derived vs. ancestral alleles (Fig. 1E,F and SOM). The two alleles act as natural controls for each other, correcting for local effects such as variation in mutation and recombination rates, or detection of singletons.

The data are normalized within 1% bins of derived allele frequency so that overall they have mean 0 and variance 1 for all frequencies from 5% —95%. By construction, positive values of SDS correspond to an increased frequency of the derived allele. In simulations, SDS follows a standard normal distribution under the null (fig. S1).

Figure 2 illustrates the properties of the SDS approach. We first investigated the relevant time-scales over which SDS might detect selection. Based on a recent model of European demography [25], we estimate that the mean tip length for a neutral sample of 3,000 individuals is 75 generations, or roughly 2,000 years (Fig. 2A). Since SDS aims to measure changes in tip lengths of the genealogy, we conjectured that it would be most likely to detect selection approximately within this timeframe.

**Figure 2.**
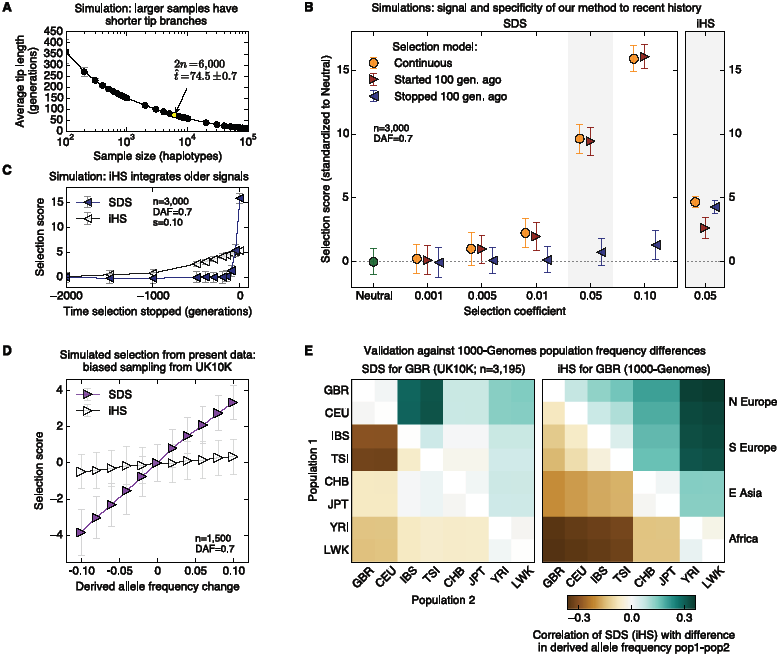
Properties of SDS. ***A.** The mean length of terminal (tip) branches as a function of sample size, for a demographic model with strong recent growth [25]. **B.** Power simulations for SDS (mean +/- SE) with 3,000 individuals under three models of selection with current derived allele frequency of 0.7: continuous hard sweep (orange); soft sweep starting 100 generations before present (red); and hard sweep that stopped 100 generations before present (blue). Right-hand panel: simulations for iHS using same models, s=0.05. **C.** Expected values of SDS and iHS, for hard sweeps that stopped in the past (s=0.10), followed by neutral drift (n=3,000). **D.** Power to detect simulated selection within UK10K data. We performed biased down-sampling of UK10K individuals so as to change the frequencies at randomly chosen SNPs. This procedure simulates an instantaneous pulse of strong selection on standing variation using real data. **E.** Allele frequency differences between extant populations vs. SDS or iHS. SDS is most correlated with the difference between southern and northern Europe, while iHS reflects longer timescales: namely Europe vs. Africa divergence. (GBR=British; CEU=Utah residents (northwest European ancestry); IBS=Iberians (Spain); TSI=Tuscans (Italy); CHB=Han (China); JPT=Japanese; YRI=Yoruba (Nigeria); LWK=Luyha (Kenya).)*

Indeed, in simulated sweep models with samples of 3,000 individuals (Fig. 2B,C and fig. S2), we find that SDS focuses specifically on very recent time scales, and has equal power for hard and soft sweeps within this timeframe. At individual loci, SDS is powered to detect ∼2% selection over 100 generations. Moreover, SDS has essentially no power to detect older selection events that stopped >100 generations before the present. In contrast, a commonly-used test for hard sweeps, iHS [12], integrates signal over much longer timescales (>1,000 generations), has no specificity to the more recent history, and has essentially no power for the soft sweep scenarios.

We next turned our attention to analysis of data from the UK10K project [26]. After removing individuals with substantial non-British ancestry or unusual numbers of singletons, we were left with 3,195 individuals with whole genome sequence data (SI and fig. S3).

At this point we wanted to validate that SDS can detect recent selection from standing variation in real data with incomplete detection of singletons. To this end, we performed repeated (biased) subsampling of 1,500 individuals from the UK10K data in such a way as to change the allele frequencies at a target SNP by 1%-10% (Fig. 2d and fig. S4). This procedure, which models strong instantaneous selection from present variation, shows that each 1% change in allele frequency results in about a 0.3-0.4 standard deviation shift in mean SDS. In contrast, iHS has essentially no power in this scenario. In summary, we expect to have power to detect very recent strong selection at individual loci, or weaker signals shared within classes of sites.

We next used the entire set of 3,195 genomes to compute SDS for 4.5 million autosomal SNPs with minor allele frequency >5% that passed our SNP-level filters. We tested whether SDS predictions are correlated with allele frequency differences between populations (Fig. 2e and fig. S5). Across all SNPs, SDS is most strongly correlated with differences between southern and northern European populations (Spearman’s ρ=0.32±0.005). In contrast, iHS measured in a British sample is most correlated with African-European differences. This provides further evidence that SDS measures historical changes in allele frequencies and reflects much more recent times than iHS.

Figure 3A summarizes the SDS signals across the genome. The single largest peak is at the lactase locus, with a maximum SDS score of 10.0 (p=1x10^−23^; Fig. 3B, fig. S8). Lactase is a well-known target of selection in Europeans [4,24], however our data show that strong selection persisted well into the last 2,000 years.

**Figure 3.**
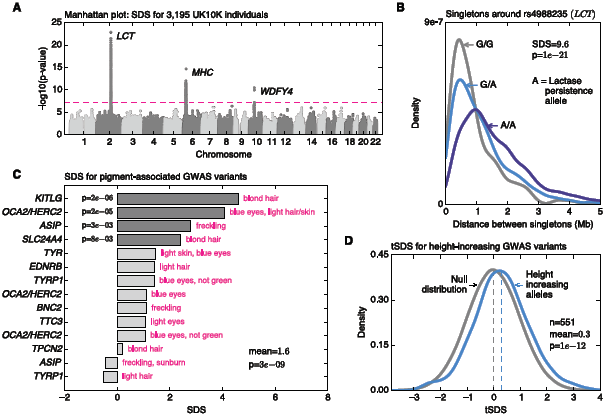
Overview of signals. ***A.** Manhattan plot of SDS p-values indicates regions of genomewide significant signals (p<5x10-8; p-values are two-sided tail probabilities of standard normal). B. Distributions of singleton distances at the lactase locus, partitioned by genotypes at the causal site. Compare to simulated signals (Fig. 1e). C. SDS signals for a curated set of segregating variants with known effects on pigmentation shows systematic increases in derived allele frequencies (one-sided p-values). **D.** Distribution of tSDS scores at 551 height-associated SNPs. tSDS scores are polarized so that tSDS>0 implies increased frequency of the “tall” allele*.

The second highest group of signals consists of multiple peaks within the extended HLA region (max SDS=7.9; p=2x10^−15^). HLA plays an essential role in immunity and has been associated with many complex traits and diseases [27]. Although we observe instances of reference bias in read mapping within this highly polymorphic region, the strong SDS signals are not an artifact of mapping bias (SOM, figs. S5c, S6). HLA is a well-known target of long-term balancing selection [28,29]; our results also imply powerful short-term directional selection on particular alleles or haplotypes. Interestingly, a recent study that used ancient DNA to study selection in Europeans during the past ∼8,000 years also reported a signal for directional selection within HLA [24]; but their top hit is in the opposite direction of the SDS score (fig. S7). This may indicate complex dynamics of selection at HLA region during recent history.

Third, SNPs in the neighborhood of *WDFY4* also cross genome-wide significance (p=3.5x10^−11^). The nature of selection is unclear, although variants in the region have previously been associated with lupus and noted for high F_ST_ values [30].

We next turned to consideration of specific variants with known effects. For GWAS-associated variants overall [31], the variance of SDS is significantly inflated (p=7x10^−16^; p=5x10^−7^ after excluding variants near HLA and lactase; SOM), indicating that GWAS variants are more likely to move up or down in frequency than random SNPs. We then tested for excess variance in SDS within categories of related variants reported in the GWAS catalog. The strongest enrichment is for variants associated with pigmentation (Fig. 3C and figs. S9, S10). While the major determinants of human skin color are near fixation and thus not testable by SDS, there is a strong overall enrichment of selection in favor of derived variants associated with lighter pigmentation, especially of hair and eye color (p=3x10^−9^ for mean SDS>0). The strongest signal is for selection in favor of a blond hair variant at the *KITLG* locus, currently at 12% frequency (p=2x10^−6^), and a weaker signal in favor of blond hair at *SLC24A4* (p=8x10^−3^). We also replicate a known signal for blue eyes at the *HERC2*/*OCA2* locus (p=2x10^−5^) [12,24]. We speculate that recent selection in favor of blond hair and blue eyes may reflect sexual selection for these phenotypes in the ancestors of the British, as opposed to the longer-term trend toward lighter skin pigmentation in non-Africans, generally thought to have been driven by the need for Vitamin D production [32].

Thus far we have described strong selection signals at individual loci. However, it has been proposed that another major mechanism of adaptation may be through polygenic selection on complex traits [16]. This mode of selection would act through small, directed shifts in allele frequencies at many loci. Polygenic selection can potentially change phenotypes rapidly, but would leave only weak signals at individual loci. The best evidence of polygenic adaptation is for height, which shows a signal of differential selection on height in northern Europeans relative to southern Europeans [17 –19,33]. A recent study of prehistoric Europeans found selection both for decreased height in southern Europeans, and a weak signal of an increase in steppe populations that contributed to northern Europeans, within the past ∼5,000 years [24].

To evaluate whether we could detect this signal using SDS, we considered a set of 551 independent height-associated SNPs from a recent meta-analysis [34] (Fig. 3D). For each SNP we modified the sign of SDS so that it reflects the change in frequency of the height-increasing allele (as opposed to derived allele for standard SDS). We refer to these modified scores as “trait-increasing SDS” (or tSDS). As shown in Figure 3D, tSDS scores for the height-associated SNPs are significantly skewed toward positive values (p=4x10^−11^), indicating that the “tall” alleles have been systematically increasing in frequency within the past ∼2,000 years in the ancestors of the British.

Aside from height and a recent report for BMI [19], evidence for selection on other complex traits has generally been weak or absent (e.g., [18,24,35]). However, it is now clear that most trait heritability is due to SNPs that do not reach genome-wide significance in association studies [36]. Indeed, recent work shows that for many traits, much of the genome is linked to variants with small, but nonzero effects [37,38]. We thus hypothesized that we could increase power to detect selection by including all SNPs, not just genome-wide significant hits. Indeed for height, the correlation between tSDS and GWAS p-value is far more significant using all SNPs that have a reported effect size, than the analysis based on significant hits alone (Spearman ρ=0.078; p=9x10^−74^; block jackknife used to correct the p-value for LD, SOM; fig. S11).

However, one concern when studying polygenic selection is that population structure might drive spurious correlations [17]: contemporary GWAS studies aim to remove the confounding effects of population structure, but it is possible that some residual effect of population structure might still remain. This in turn could correlate with SDS (Fig. 2E). To solve this problem, some studies have used family-based association data [17,19], which provide stringent control for structure, albeit with much smaller available sample sizes.

We thus analyzed a recent family-based GWAS for height [19]. The correlation between SDS and GWAS effect size is even stronger in these data than in the much larger meta-analysis (Spearman ρ=0.094; p=9x10^−163^). We speculate that the published height meta-analysis may actually over-correct for population structure since the structure is pervasively correlated with phenotypic signal.

Remarkably, mean tSDS is positive across nearly the entire range of p-values. It may seem counter-intuitive that, in aggregate, even non-significant SNPs could have a detectable association with tSDS. However, a recent method (ashR [39]) estimates that 85% of SNPs in this data set are associated with a non-zero effect on height (including indirect effects through tagging of LD partners), and that the direction of effect is estimated correctly for 68% of all SNPs (fig. S12). In summary, our results indicate that polygenic selection on height has affected allele frequencies across most of the genome.

We expanded our test to consider a total of 43 traits for which genome-wide GWAS data are available (SOM, table S1). Unfortunately, large-scale family studies are not available for most traits. As an alternative, we used LD Score regression to verify correlations between traits and SDS [37,40]. LD Score regression tests whether the covariance between two genome-wide association studies increases with the amount of LD. Such a signal is expected if two highly polygenic signals are truly correlated, but there should be little, or no, such signal if an association is driven purely by population structure. Here we adapted this method to test whether the covariance between SDS and GWAS effect sizes increases with LD as would be expected for a signal of polygenic selection (SOM). Indeed for height, there is a strong LD Score signal of selection, both in the family data (p=3x10^−17^) and the meta-analysis data (p=2x10^−11^, Fig. 4b, fig. S11).

**Figure 4.**
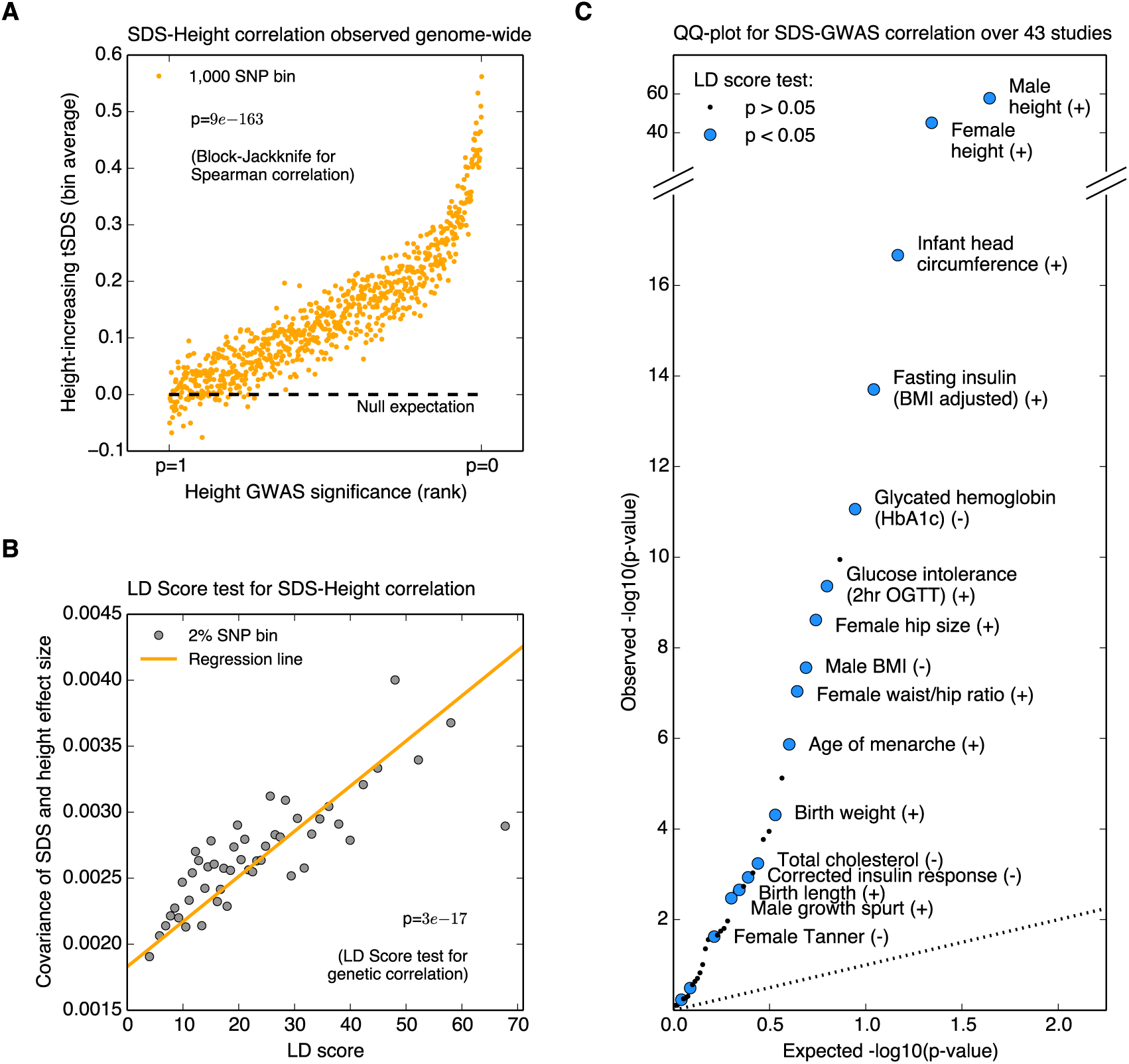
Signals of polygenic adaptation. ***A.** Mean tSDS of SNPs, where tSDS is polarized according to the estimated direction of effect of each SNP on height in a recent family-based study [19]. The x-axis is ordered from least significant SNPs (p∼1) to most significant (p∼0) and SNPs are placed into bins of 1,000 consecutive SNPs for easier visualization. **B.** Covariance of height effect size and SDS, as a function of LD score, indicates that selection on height is truly polygenic (LD score p=2x10^−11^). **C.** QQ-plot testing for a correlation between GWAS effect size and tSDS for 43 traits. Significant traits that are also nominally significant by the LD Score method are colored blue and labeled (p<0.05, LD Score one-sided test)*.

Strikingly, many traits show highly significant associations between SDS and GWAS effect size (Fig. 4C). Most of the significant traits also show nominally significant associations in a concordant direction by the more stringent LD Score test (see labeled traits in Fig. 4C). While height has the strongest signal, we also see signals for a variety of other traits. These include additional anthropometric traits such as selection toward increased infant head circumference and birth weight, and increases in female hip size; as well as selection on variants underlying metabolic traits including levels of insulin and glycated hemoglobin. Some of the results hint at sexually dimorphic adaptations: in addition to the female-specific signal for increased hip size, we observe a male-specific signal for decreased BMI; we further see a signal in favor of later sexual maturation in women, but not in men. While many of these signals are highly intriguing, and some match known phenotypes of modern British (SOM), it remains to fully determine the confounding role—if any—of population structure in contributing to these signals.

In summary, we have introduced a powerful new method for inferring very recent changes in allele frequencies. The method will be widely applicable across human populations and other species. We have shown that selection on milk digestion, and on physical attributes including blond hair and blue eyes continued well into historical times. Our results also suggest that polygenic selection played an important role in the evolution of many complex traits. Selection on height alone has affected allele frequencies at most SNPs genome-wide, and we anticipate that competing effects of linked SNPs on different traits may be an important feature of short-term adaptation. It is important to note that the measured traits are, themselves, not necessarily direct targets of selection; it is likely that each of these traits shares variants with other correlated phenotypes as well [40]. Nonetheless, our results suggest that selection on complex traits has been an important force in shaping both genotypic and phenotypic variation within historical times.

## Acknowledgments

We thank H. Fraser, J. Pickrell, M. Przeworski, G. Sella and members of the Pritchard lab for comments. This work was supported by NIH grants ES025009, 5T32HG000044-19 and MH101825, and by the Howard Hughes Medical Institute.

